# Quantification of malaria antigens PfHRP2 and pLDH by quantitative suspension array technology in whole blood, dried blood spot and plasma

**DOI:** 10.1101/730499

**Authors:** Xavier Martiáñez-Vendrell, Alfons Jiménez, Ana Vásquez, Ana Campillo, Sandra Incardona, Raquel González, Dionicia Gamboa, Katherine Torres, Wellington Oyibo, Babacar Faye, Eusebio Macete, Clara Menéndez, Xavier C. Ding, Alfredo Mayor

## Abstract

**Background:** Malaria diagnostics by rapid diagnostic tests (RDTs) relies primarily on the qualitative detection of *Plasmodium falciparum* histidine-rich protein 2 (PfHRP2) and *Plasmodium sp* lactate dehydrogenase (pLDH). As novel RDTs with increased sensitivity are being developed and implemented as point of care diagnostics, highly sensitive laboratory based assays are needed for evaluating RDTs performance. Here, a quantitative suspension array technology (qSAT) was developed, validated and applied for the simultaneous detection of PfHRP2 and pLDH in a variety of clinical samples (whole blood, plasma and dried blood spots) from different endemic countries.

**Results:** The qSAT was specific for the target antigens, with analytical ranges of 6.8 to 762.8 pg/ml for PfHRP2 and 78.1 to 17076.6 pg/ml for *P. falciparum* (Pf-LDH). The assay detected *P. vivax* LDH (Pv-LDH) at a lower sensitivity than Pf*-*LDH (analytical range of 1093.20 to 187288.5 pg/ml). Both PfHRP2 and pLDH levels determined using the qSAT showed to positively correlate with parasite densities determined by quantitative PCR (Spearman r=0.59 and 0.75, respectively) as well as microscopy (Spearman r=0.40 and 0.75, respectively), suggesting the assay to be a good predictor of parasite density.

**Conclusion:** This immunoassay can be used as a reference test for the detection and quantification of PfHRP2 and pLDH, and could serve for external validation of RDTs performance, to determine antigen persistence after parasite clearance, as well as a complementary tool to assess malaria burden in endemic settings.

## INTRODUCTION

The availability of field-deployable malaria rapid diagnostic tests (RDTs) in recent years has markedly facilitated access to malaria diagnostics. Since the World Health Organization (WHO) recommendations in 2010 to test all suspected malaria cases (1), RDTs have gained a crucial role in the management of malaria clinical cases, as well as for malaria surveillance. Malaria RDTs have supplanted conventional light microscopy in many endemic areas as the standard of practice, accounting in 2017 for 75% of all diagnostic tests done in sub-Saharian Africa, were most RDTs are distributed (66%) (2). The vast majority of RDTs used worldwide are based on the detection of parasite bioproduct histidine-rich protein 2 (PfHRP2), expressed only in *Plasmodium falciparum*, and the parasite metabolic enzyme lactate dehydrogenase (pLDH), present in all human-infecting *Plasmodium* species.

PfHRP2 is a water-soluble glycoprotein produced by the parasite throughout its asexual lifecycle and early sexual stages; it is expressed on the surface of infected erythrocytes and released into the peripheral blood circulation during schizogony (3,4). Given the ability of mature *P. falciparum* parasites to sequester in vascular beds during the last half of their asexual life-cycle, where they are not accessible for microscopic diagnosis, it has been proposed that the quantitative detection of PfHRP2 can provide a more accurate measurement of parasite biomass and potentially assist in determining the prognosis of severe malaria (5–7). During pregnancy, *P. falciparum* infections can remain undetectable in peripheral blood as the parasites sequester in the intervillous spaces of the placenta by specific adhesion to chondroitin sulphate A (8,9). In such scenario, PfHRP2-detecting RDTs have showed to have higher sensitivity on peripheral blood compared to conventional light microscopy (10), although still lower than PCR (11).

RDTs detecting PfHRP2 only are the most widely used products (12), accounting for 66% of the 276 million RDTs sold worldwide in 2017 (2). Nonetheless, PfHRP2-detecting RDTs have been suggested to have limited clinical specificity for diagnosis of current malaria infection in areas of high transmission (13) and following treatment (14,15) due to the persistence of the protein in the blood circulation after parasite clearance. The time span of a positive test result following parasite clearance is mainly dependent on the duration and density of parasitaemia prior to treatment, with values ranging from 26 days in Ugandan children with parasitaemia less than 1,000 parasites per microliter (p/μl) up to 37 days for parasite density >50,000 p/μl (16).

The parasite LDH is a metabolic enzyme required for survival and is produced by all five *Plasmodium* species infective to humans (17,18). In contrast to PfHRP2, pLDH does not persist in blood after clearance of malaria infections and is therefore a better marker of acute and current infection (19). Upon treatment, pLDH clearance in blood has been shown to closely track with that of parasites, suggesting pLDH to be a suitable predictor for treatment failure (20). However, sensitivity of RDTs based on this antigen is generally lower than that of PfHRP2-based RDTs (21).

Currently, enzyme-linked immunoabsorbent assay (ELISA) is the standard of practice immunoassay for the detection and quantification of PfHRP2 and pLDH, and is used as an external validation tool for RDTs performance. ELISAs are however costly, time and sample consuming, and generally only allow for the detection of one analyte at the time. The recent release of a highly-sensitive RDT for PfHRP2 (Alere™ Malaria Ag P.f), with two to ten-fold higher sensitivity than other currently available RDTs (22,23), as well as the work in progress to develop new generation pLDH-based RDTs, underpin the need for new highly-sensitive laboratory based reference immunoassays than can provide lower limit of detection than classical ELISAs (24–28). Highly sensitive quantitative assays should not only be a more suitable tool for validation of new-generation RDTs, but could also be used to better understand antigen kinetics, particularly that of PfHRP2, and to support malaria surveillance. In this work, we present a high-throughput quantitative suspension array approach based on the Luminex technology that allows for the simultaneous and highly sensitive detection and quantification of PfHRP2 and pLDH antigens in different biological samples (whole blood, plasma, and dried blood spots). This assay provides an additional tool to externally evaluate the performance of new generation antigen detecting malaria RDTs, and can be used for research purposes to address biological questions such as PfHRP2 persistence and the relationship between antigen levels and disease severity.

## MATERIALS AND METHODS

### Development and optimization of the bead suspension array

#### Biotinylation of detection mAbs

Detection monoclonal mouse IgG α-PfHRP2 (MBS834434, MyBioSource, San Diego, CL) and monoclonal mouse IgG α-PAN-pLDH (PA-2, AccessBio, Somerset, NJ) were biotinylated using the EZ-Link Sulfo-NHS-Biotin Kit (21435, Thermo Fisher Scientific, Waltham, MA) according to the manufacturer’s instructions with minor modifications (See Additional file 1: Text1).

#### Coupling of mAbs to magnetic beads

Coupling of magnetic microspheres was performed similarly as described elsewhere (29). Briefly, two MagPlex® microspheres (Luminex Corp., Austin, Texas) with different spectral signatures selected for the detection of PfHRP2 and PAN-pLDH were washed with distilled water and activated with Sulfo-NHS (N-hydroxysulfosuccinimide) and EDC (1-ethyl-3-[3-dimethylaminopropyl] carbodiimide hydrochloride) (Pierce, Thermo Fisher Scientific Inc., Rockford, IL), both at 50 mg/mL, in activation buffer (100 mM Monobasic Sodium Phosphate, pH = 6.2). Microspheres were washed with 50 mM MES potassium salt (4-morpholineethane sulfonic acid, Sigma Aldrich, St. Louis, MO) pH 5.0 to 10000 beads/µl, and covalently coupled with capture antibodies against PfHRP2 (MBS832975, MyBiSource, San Diego, CL) and PAN-pLDH (PA-12, AccessBio, Somerset, NJ), both at a concentration of 25 µg/ml. Beads were incubated on a rotatory shaker overnight at 4 °C and protected from light. Microspheres were blocked with PBS-BN (PBS with 1% BSA and 0.05% sodium azide (Sigma, Tres Cantos, Spain), and re-suspended in PBS-BN (from now on named assay buffer) to be quantified on a Guava PCA desktop cytometer (Guava, Hayward, CA) to determine the percentage recovery after the coupling procedure. Coupling validation was performed by incubating 50 µl of each bead suspension (2000 beads/well) with 50 µl α-mouse IgG-Biotin (B7401-1ML, goat anti-Mouse IgG-Biotin, Sigma Aldrich, St. Louis, MO) at 1:1000 dilution in a 96-well flat bottom plate for 2 hours in gentle agitation. The plate was washed by pelleting microspheres using a magnetic separator (40-285, EMDMillipore, Burlington, MA) and re-suspended with wash buffer (0.05% Tween 20/PBS). Beads were incubated with 100 µl of streptavidin-phycoerythrin (42250-1ML, Sigma Aldrich, St. Louis, MO) diluted 1:1000 in assay buffer for 30 minutes in gentle agitation in the dark. Finally, the beads were washed and re-suspended in assay buffer, and the plate was read using the Luminex xMAP® 100/200 analyser (Luminex Corp., Austin, TX). A reading higher than 25000 median fluorescence intensity (MFI) implied a successful coupling reaction. Coupled beads were stored multiplexed at a concentration of 1000 beads/µl/region at 4 °C and protected from light.

To optimize the coupling concentration of detection antibodies, a concentration range from 10 to 100 µg/ml of α-PfHRP2 and α-PAN-pLDH monoclonal antibodies (mAbs) was conjugated to magnetic beads, and assayed against serially diluted recombinant PfHRP2 and pLDH and a selection of plasma samples from *P. falciparum* positive individuals. The mAb concentration that provided the highest the MFI values was selected as the optimal concentration.

#### PfHRP2 and pLDH reference materials

Recombinant PfHRP2 protein type A from FCQ79 *P. falciparum* strain expressed in *Escherichia coli* (890015, Microcoat GmbH, Germany) was selected as PfHRP2 reference material. Antigen concentration after reconstitution was determined by ELISA (Malaria Ag CELISA, CeLLabs). Purified recombinant *P. falciparum* and *P. vivax* pLDH proteins expressed in insect cells (3001, ReliaTech GmbH, Germany) were used as reference material. The pLDH concentrations were measured in a previous study using a commercially available ELISA (QUALISA Malaria kit, Qualpro Diagnostics, India) (21). Reference materials were used to prepare the standard curves for the bead suspension array, starting at concentrations of 50 ng/ml for PfHRP2 type A and at 1000 ng/ml for *P. falciparum* and *P. vivax* pLDH. The WHO International Standard for *Plasmodium falciparum* antigens was provided by the National Institute for Biological Standards and Control (Ridge, UK) (NIBSC code: 16/376, (30)). The WHO International Standard for *P. falciparum* antigens was quantified, and the obtained antigen concentrations in pg/ml were used to calculate the number of antigen picograms corresponding to 1 International Unit (IU).

#### Optimization of assay standard curves

Standard curves were prepared for the detection of PfHRP2 and pLDH. The conjugated beads were incubated with serial dilutions of recombinant PfHRP2 types A, B and C and recombinant *Pf* and *Pv* pLDH in assay buffer to produce standard curves ranging from 50,000 to 0.024 pg/ml for PfHRP2, and from 1,000,000 to 0.48 for both *P. falciparum* pLDH and *Pv* pLDH (Figure 1B) (for a more detailed assay procedure, see Additional file 1: Text2).

**Figure 1.**
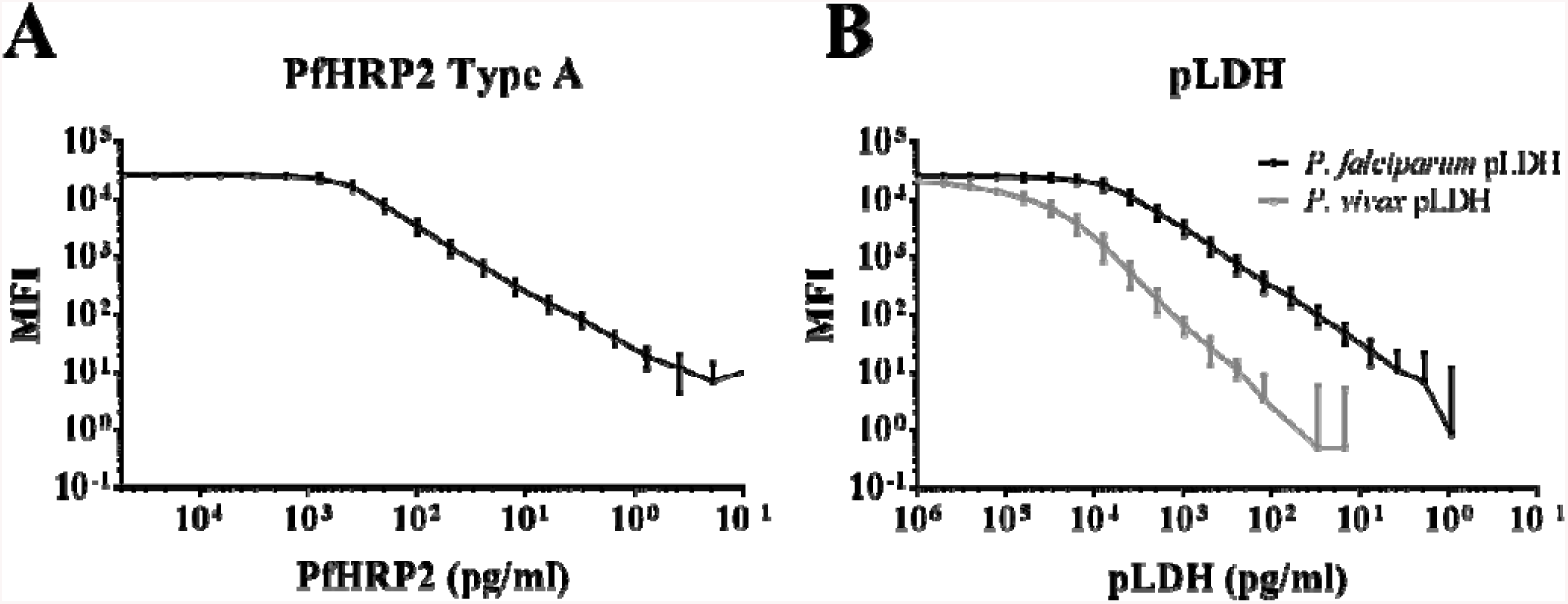
Calibration curves to detect PfHRP2, *Pf* pLDH and *Pv* pLDH. Recombinant *P. falciparum* (*Pf*) HRP2 type A (A) and *Pf* (B, back line) and *P. vivax* (*Pv)* (B, grey line) pLDH were serially diluted to investigate the assay analytical range. Error bars show the standard deviation of the mean from 66 independent reads for PfHRP2 type A and *Pf* pLDH, and 12 reads for *Pv* pLDH. X axis: MFI value after subtraction of the background.

### Assay parameters

#### Limit of detection, limits of quantification and range

A calibration curve prepared with serially diluted reference PfHRP2 and *P. falciparum* pLDH was assayed in 66 runs on the Luminex xMAP® 100/200 analyser, along with 2 blank samples (consisting of assay buffer alone) per run. For *Pv* pLDH, serial dilutions of reference antigen were assayed in 6 independent runs. The lower limits of detection (LLOD), defined as lowest amount of analyte which can be detected, and of quantification (LLOQ), defined as the lowest concentration of an analyte in a sample that can be quantified, were determined by measuring the MFI of 132 wells containing blank samples. The upper limit of quantification (ULOQ), corresponding to the highest concentration that can be quantitatively determined, was defined as the maximum value of the fitted mean standard curve minus its 10% to avoid quantifying samples falling close to the saturation plateau. The analytical range was set within the lower and the upper limits of quantification.

To quantify the LLOD and the LLOQ, 3 and 6 standard deviations (SD) were added to the mean MFI of blanks (n = 132), respectively. Each calibration or standard curve was fitted using a five parameters logistic (5PL) regression, and the mean curve was calculated. To present the LLOD and the LLOQ as concentration values, the calculated MFI values were interpolated to the mean calibration curve.

#### Dilution linearity and accuracy

Dilution linearity and accuracy were evaluated on the same serial dilutions of recombinant PfHRP2 type A and *P. falciparum* pLDH read over 66 independent runs. Dilution linearity was calculated as the mean percent change in dilution-corrected concentration from one dilution to the previous one within the assay range. Dilution linearity was considered acceptable if the percent change in concentration did not exceed the recovery range of 80-120%. Accuracy was determined as the mean percent deviation (% DEV) from the expected concentration, calculated by diving the difference between the experimental value and the expected value and then multiplying by 100. Acceptable accuracy was defined as the %DEV not surpassing by 20% the expected concentration (by 25% for samples with concentrations falling at the LLOQ and ULOQ).

#### Precision

Intra-assay and inter-assay precision were evaluated by assaying cultured *P. falciparum* W2 strain spiked in assay buffer at five dilutions spanning a wide range of antigen concentration in triplicate over four runs. Intra-assay precision over the four runs was defined as the average coefficient of variation (% CV) of individual samples. The % CV for each sample was calculated by determining the standard deviation (SD) of the three replicate results, dividing it by the mean of the triplicate results, and multiplying by 100. Inter-assay precision was defined as the overall % CV, calculated by dividing the SD of plate means by the mean of plate means and then multiplying by 100. Calculations were performed on non-transformed MFI values. Precision was considered acceptable when % CV did not exceed 10% for intra-assay variation and 20% for inter-assay variation (30).

#### Selectivity

To investigate the selectivity of the assay for the target antigens, 75 plasma samples from 25 Spanish pregnant women never exposed to malaria were assayed to demonstrate that the bead suspension array does not detect plasma components other than the target antigens (PfHRP2 and pLDH).

### Study samples

To test against samples collected in endemic areas, different sample sets were assayed (characteristics of clinical samples used are summarized in Table 1).

**Table 1.**
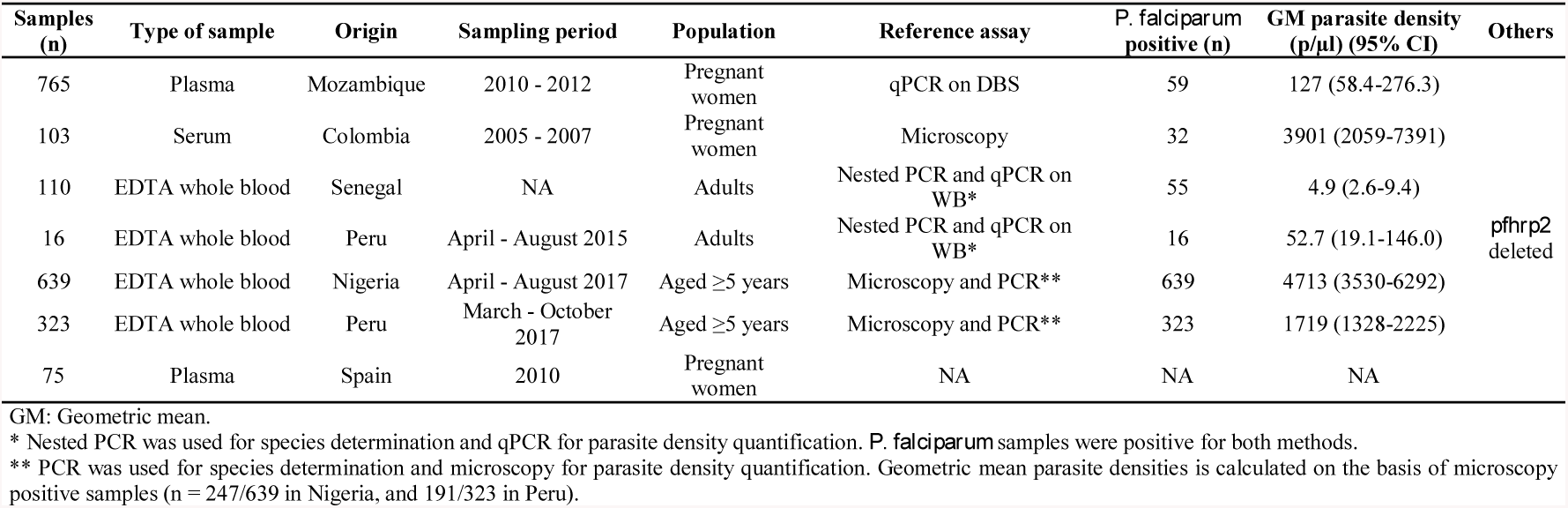
Clinical samples tested on the qSAT assay.

| Samples (n) | Type of sample | Origin | Sampling period | Population | Reference assay | *P. falciparum* positive (n) | GM parasite density (p/μl) (95% CI) | Others |
| --- | --- | --- | --- | --- | --- | --- | --- | --- |
| 765 | Plasma | Mozambique | 2010 - 2012 | Pregnant women | qPCR on DBS | 59 | 127 (58.4-276.3) |  |
| 103 | Serum | Colombia | 2005 - 2007 | Pregnant women | Microscopy | 32 | 3901 (2059-7391) |  |
| 110 | EDTA whole blood | Senegal | NA | Adults | Nested PCR and qPCR on WB* | 55 | 4.9 (2.6-9.4) |  |
| 16 | EDTA whole blood | Peru | April - August 2015 | Adults | Nested PCR and qPCR on WB* | 16 | 52.7 (19.1-146.0) | *pfhrp2* deleted |
| 639 | EDTA whole blood | Nigeria | April - August 2017 | Aged ≥5 years | Microscopy and PCR** | 639 | 4713 (3530-6292) |  |
| 323 | EDTA whole blood | Peru | March - October 2017 | Aged ≥5 years | Microscopy and PCR** | 323 | 1719 (1328-2225) |  |
| 75 | Plasma | Spain | 2010 | Pregnant women | NA | NA | NA |  |

GM: Geometric mean.
\* Nested PCR was used for species determination and qPCR for parasite density quantification. *P. falciparum* samples were positive for both methods.
\*\* PCR was used for species determination and microscopy for parasite density quantification. Geometric mean parasite densities is calculated on the basis of microscopy positive samples (n = 247/639 in Nigeria, and 191/323 in Peru).

#### *Plasmodium falciparum* culture samples and *P. vivax* clinical samples

W2, Benin I, Borneo and Santa Lucia *P. falciparum* strains were cultured under standard hypoxic conditions. Culture in exponential growth phase was harvested, infected red blood cells were spun down, aliquoted, and frozen at −80 °C as previously described (21). *Plasmodium vivax* isolates were collected from symptomatic adult volunteers with a *P. vivax* mono-species infection as confirmed by microscopy during a specimen collection campaign organized in April 2016 in the area of Iquitos (Peru).

#### Plasma and serum samples

PfHRP2 and pLDH were measured in 765 plasma samples collected at 3 time points during pregnancy from 255 pregnant women residing in Manhiça (Southern Mozambique) who participated in a clinical trial of intermittent preventive treatment during pregnancy (IPTp) from 2010 to 2012 (31,32), and in 103 serum samples from 77 pregnant women in the Urabá-Antioquia region (Colombia) collected between 2005 and 2007 (33). Additionally, 75 plasma samples collected at 3 time points from 25 pregnant women never exposed to malaria, who attended the Hospital Clínic of Barcelona during pregnancy and delivery in 2010, were included in the assay as negative controls. Plasma and serum samples were stored at −80 °C. Infection status and parasite densities were previously determined by qPCR on DBS for samples from Mozambique (34), and by light microscopy (LM) in Colombian samples.

#### Whole blood samples

EDTA-anticoagulated whole blood samples were collected from consenting asymptomatic adults with no recent clinical episode of malaria (past four weeks) during cross-sectional surveys in Peru (35), and Senegal. Samples were assessed and categorized as *P. falciparum* mono-species infection or *Plasmodium* negative samples using nested PCR, and parasitaemia was quantified using quantitative PCR as described previously at the Hospital for Tropical Diseases (UK) (36). The *pfhrp2* gene status of *P. falciparum* PCR positive samples was investigated by PCR as previously described (37). Whole blood samples from asymptomatic adults were used to prepare dried blood spots (DBS) (See Additional file 1: Text4). EDTA-anticoagulated whole blood samples were collected between March and October 2017 in Peru Amazon region and Nigeria Lagos state from consenting symptomatic (with fever within the last 3 days) and asymptomatic (no fever history in past 3 days) patients enrolled during a clinical trial of a new multiplex fever diagnostic test. Antigens were quantified in those samples that were positive for *P. falciparum* by PCR (n= 323 in Peru and 629 in Nigeria). Individuals participating in this clinical trial had been previously tested by on-site microscopy (final result based on reading from 2 independent microscopists), and by SD BIOLINE Malaria Ag P.f (HRP2/pLDH) (05FK90, Abbott, Chicago, IL) in Nigeria and by CareStart™ Malaria Pf/PAN (HRP2/pLDH) (RMRM-02571) and Carestart Pf/PAN (pLDH) Ag (RMLM-02571) (AccessBio, Somerset, NJ) RDTs in Peru.

### Statistical analysis

The relationship between the MFIs in singleplex and multiplex assays and the correlation between parasite densities and antigen levels were assessed by the non-parametric Spearman’s rank correlation method. Statistical analyses were performed with GraphPad Prism (version 6, Graphpad, Inc). The 95% confidence intervals (CI 95%) for sensitivity and specificity were calculated by Wilson score method in Microsoft Excel (2013).

## RESULTS

### Development of the bead suspension array for PfHRP2 and pLDH detection

#### Optimization of standard curves for the detection of PfHRP2 and pLDH

The coupling conditions were optimized based on a concentration range of 10 to 100 ug/mL of coupled HRP2 and pan-pLDH antibodies and testing with recombinant antigens as well as plasma samples from *Pf* infected pregnant women, showing slightly higher MFI values at 25 ug/mL (data not shown). A range of in-house biotinylated detection mAbs was tested, and the optimal concentration was found to be 1 μg/ml for the detection of both antigens (data not shown).

PfHRP2 type A, slightly higher MFI values were obtained for type A compared to types B and C (See Additional file 2: Figure S1A), similarly to previously reported data (24,25). PfHRP2 type A was selected as reference material. Recombinant *P. falciparum* pLDH was detected down to lower concentrations compared to *Pv* pLDH, indicating higher assay sensitivity for the detection of recombinant *P. falciparum* pLDH (Figure 1B). Similarly, the assay was able to detect lower concentrations of native *P. falciparum* pLDH compared to *P. vivax* pLDH (See Additional file 2: Figure S1B). Additionally, the detection of PfHRP2 and pLDH in assay buffer spiked with recombinant proteins, cultured parasites or plasma samples yielded similar MFI values in singleplex and multiplex (See Additional file 2: Figure S1C), with a clear correlation for both PfHRP2 (n = 25, r = 0.995; p < 0.001) and pLDH (n = 31, r = 0.992; p < 0.001), indicating no cross-reactivity between PfHRP2 and pLDH components.

#### Correspondence to International Units

In the qSAT assay presented here, 1 IU PfHRP2 corresponds to 23.5 pg PfHRP2, whereas 1 IU pLDH corresponds to 160 pg/ml pLDH.

### Assay parameters

#### Limit of detection, limits of quantification and range

The lower limit of detection (LLOD) of the assay was determined to be 6.0, 56.1 and 1093.20 pg/ml for recombinant PfHRP2 type A, *P. falciparum* pLDH and *P. vivax* pLDH respectively; and the lower limit of the quantification (LLOQ) was estimated at 6.8 pg/ml for PfHRP2, 78.1 pg/ml for *P. falciparum* pLDH and 1343.5 pg/ml for *P. vivax* pLDH. The ULOQ was found to be 762.8 pg/ml, 17076.6 pg/ml and 187288.5 pg/ml for PfHRP2, *P. falciparum* pLDH and *P. vivax* pLDH, respectively. The limits of detection for PfHRP2 types B and C were 17.2 pg/ml and 15.8 pg/ml, respectively.

#### Dilution linearity and accuracy

The mean percent change in dilution-corrected concentration between contiguous dilutions was 13.6 and 11.1% for PfHRP2 and *P. falciparum* pLDH, respectively, as determined over 66 independent runs. These data are within the acceptance criteria of +/-20% (38). However, at concentrations close to the ULOQ, the percent change showed an overestimation greater than 20% for both PfHRP2 and *P. falciparum* pLDH (Table 2). The overall percent deviation between the experimental concentration and the expected concentration for each serial dilution point falling within or close to the analytical range was 19.6 and 16.4% for PfHRP2 and pLDH, respectively. At concentrations falling at the LLOQ and the ULOQ, accuracy decreased both for PfHRP2 and *P. falciparum* pLDH detection as shown in Table 2.

**Table 2.** Dilution linearity and accuracy of qSAT assay.

| *Pf* pLDH |  |  |  | PfHRP2 |  |  |
| --- | --- | --- | --- | --- | --- | --- |
| Sample | Expected concentration (pg/ml) | % Change | % Deviation | Expected concentration (pg/ml) | % Change | % Deviation |
| 1 | 15625 | 41.6 | 24.2 | 781.3 | 32.8 | 23.4 |
| 2 | 7812.5 | 18.3 | 17.4 | 390.6 | 18.5 | 10.8 |
| 3 | 3906.3 | 0.9 | 17.1 | 195.3 | 6.8 | 10.2 |
| 4 | 1953.1 | 2 | 10.7 | 97.7 | 0 | 8.5 |
| 5 | 976.6 | 5.6 | 10.4 | 48.8 | 5.2 | 10.1 |
| 6 | 488.3 | 4.2 | 14.2 | 24.4 | 14.7 | 19 |
| 7 | 244.1 | 5.5 | 14.9 | 12.2 | 12.7 | 27.9 |
| 8 | 122.1 | 10.4 | 22.2 | 6.1 | 18.5 | 47 |
| Overall | - | 11.1 | 16.4 | - | 13.6 | 19.6 |

#### Precision

Intra-assay variation was 8.3% and 9.8% for PfHRP2 and pLDH, respectively. The inter-assay %CV was 8.4% for the detection of PfHRP2 and 11.2% for the detection of pLDH. For both antigens, intra-assay and inter-assay variation fell within the acceptance criteria of 15% and of 20 % variation, respectively (30).

### PfHRP2 and pLDH recovery from dried blood spots

To determine the loss of antigen when recovering PfHRP2 and pLDH from filter papers as compared to same volumes of whole blood samples, DBS were prepared with whole blood samples from Senegalese and Peruvian asymptomatic individuals (See Table 1). Blood was eluted from DBS in assay buffer (See Additional file 1: Text4) and eluted samples were assayed on the bead-suspension array along with the original whole blood samples used to prepare the DBS. The geometric mean antigen concentration obtained from DBS eluted product was 0.04 ng/ml (95% CI 0.03-0.07 ng/ml) for pfHRP2 and 0.10 ng/ml (95% CI 0.06-0.16 ng/ml) for pLDH. These concentrations are 22.8 (n=38, 95% CI 15.6-33.5) and 59.7 (n=18, 95% CI 35.4-100.6) times lower than the concentrations obtained in whole blood for PfHRP2 and pLDH, respectively (0.77 ng/ml (95% CI 0.37-1.61 ng/ml) for PfHRP2 and 5.77 ng/ml (95% CI 2.45-13.57 ng/ml) for pLDH), for identical blood volumes.

### Assay selectivity for the target antigens

An important step in the development of the assay was to investigate whether it was selective for the target antigens. Significant MFI signal for PfHRP2 and pLDH was observed in *P. falciparum* positive samples (PfHRP2: Mean = 10195, SD = 12545; pLDH: Mean = 9634, SD = 11765), whereas positive *P. vivax* samples (n=12) only showed fluorescence signal for pLDH (Mean = 12960; SD = 3735), and not for PfHRP2 (Mean = 75.0, SD = 39.3) as expected (Figure 2). Five out of 71 and seven out of 738 negative samples by microscopy and PCR, respectively, showed MFI values above the LLOQ for both PfHRP2 and pLDH, and four other *P. falciparum* positive samples by microscopy and four *P. falciparum* positive samples by qPCR yielded greater MFI values than the LLOQ for pLDH and PfHRP2 respectively. In addition, two *P. falciparum* positive samples with *pfhrp2* deletion showed MFI values above the LLOQ. Finally, all plasma samples (n=75) from Spanish malaria naïve pregnant women yielded negligible fluorescence signals for both antigens (Figure 2).

**Figure 2.**
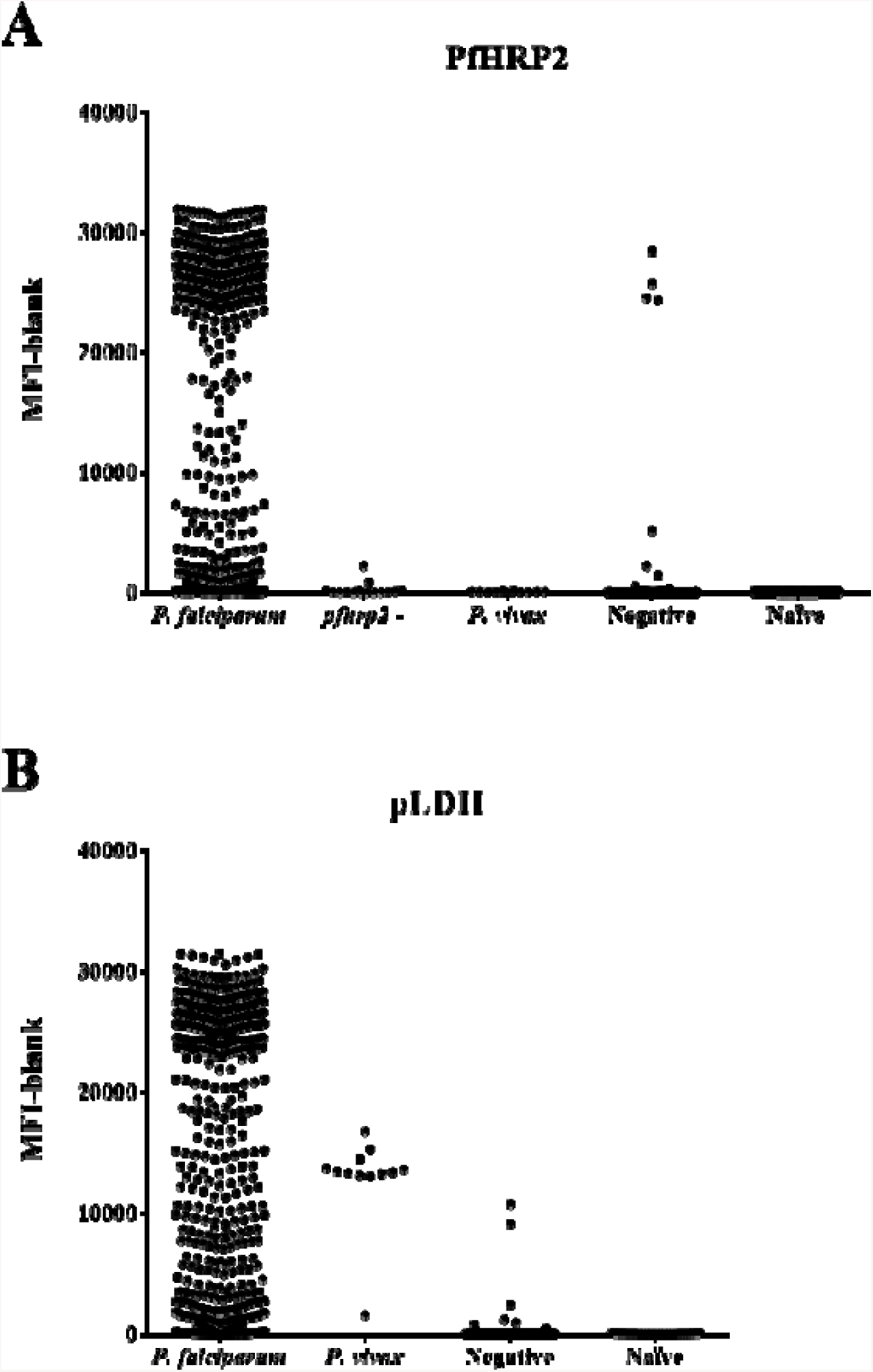
The quantitative bead suspension array is selective for PfHRP2 and pLDH. Median fluorescence intensity with blank subtracted for PfHRP2 **(A)** and pLDH **(B)** for *P. falciparum* positive samples (n = 1098), *P. falciparum* with *hrp2* gene deletions (n =16), *P. vivax* positive samples (n = 12), *Plasmodium* negative samples, and samples from naïve individuals (n =75). *pfhrp2* -: *Plasmodium falciparum* with *hrp2* gene deletion.

### Correlation between antigen levels and parasite densities

In samples positive for one or two antigens, the correlation between antigen concentrations and parasite densities was investigated. Overall, a significant correlation between PfHRP2 and parasite densities was found regardless of whether parasite densities were quantified by qPCR (Spearman r = 0.59; p < 0.0001) or microscopy (Spearman r = 0.40; p < 0.0001) (Figure 3). pLDH levels showed a higher correlation with parasite densities compared to PfHRP2, both in samples for which densities were determined by qPCR (Spearman r = 0.75; p < 0.0001) and by microscopy (Spearman r = 0.75; p < 0.0001) (Figure 3). The correlation between parasite densities and antigen levels differed across the different sample sets analysed (See Additional file 3: Table S1). Interestingly, the correlation between pLDH levels with parasite densities in whole blood samples from Peru (Spearman r = 0.76; p < 0.0001) and Nigeria (Spearman r = 0.78; p < 0.0001) was very similar, whereas for PfHRP2, a better correlation with parasite densities was found in samples from Nigeria (Spearman r = 0.47; p < 0.0001) compared to samples from Peru (Spearman r = 0.20; p = 0.0308).

**Figure 3.**
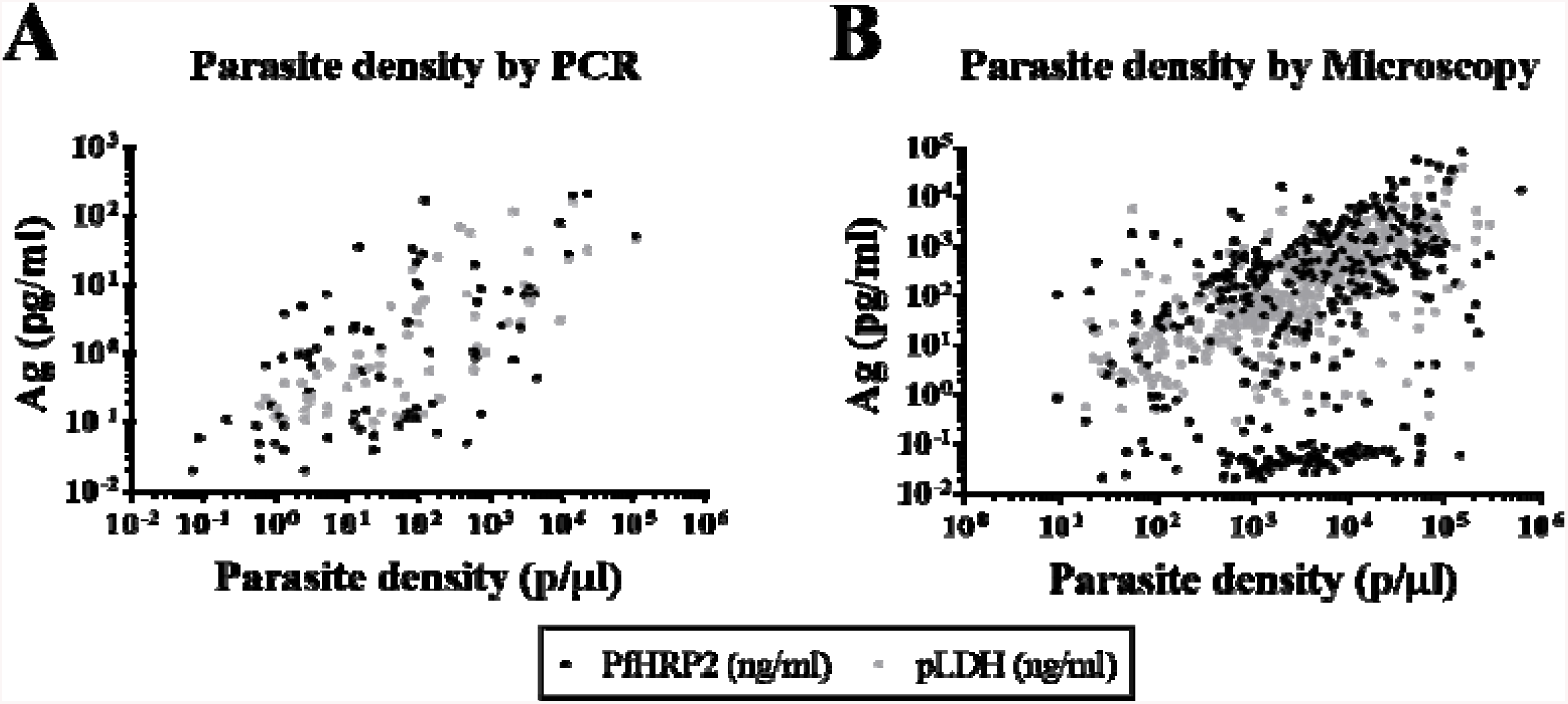
Antigen levels correlate with parasite densities. Correlation of parasite densities (p/µl) with PfHRP2 and pLDH concentration (pg/ml) in *P. falciparum* positive samples by PCR (**A**), and by microscopy (**B**).

## DISCUSSION

In the present study, we have established a quantitative suspension array based on Luminex technology for the simultaneous detection and quantification of *P. falciparum* HRP2 and *P. falciparum* and *P. vivax* pLDH, which allows to determine protein concentrations as low as 6.0, 56.1 and 1042.7 pg/ml, respectively. Hence, the assay provides increased sensitivity compared to commercially available ELISA kits which have LODs of approximately 400 pg/ml and 1000 pg/ml for PfHRP2 and pLDH, respectively (27,39). The assay shows good levels of dilution linearity, accuracy and precision, and can be used to effectively and rapidly quantify malaria antigens in large quantities of different biosamples.

The performance of the bead suspension array to quantify PfHRP2 and pLDH was evaluated using reference recombinant proteins as well as cultured parasites, and in different biofluids from malaria exposed and malaria naïve individuals. The assay is selective for the target antigens and has an analytical range of 6.8 to 762.8 and of 78.1 to 17076.6 pg/ml for PfHRP2 and *P. falciparum* pLDH, respectively. Additionally, the assay can also quantify *Pv* pLDH down to 1211.6 pg/ml. The assay analytical sensitivity to detect PfHRP2 is comparable to that of a recently developed bead suspension assay based on Luminex technology (25), as well as to other immunoassays that use different technologies (20,28). This suggests that with the current technology available for the quantification of PfHRP2 using antibodies, the lowest limit of detection achievable is in the range of 0.5 to 10 pg/ml. The limit of detection for pLDH is more divergent across assays, ranging from approximately 10 pg/ml (28) up to 4000 pg/ml (25), but in all assays it is always higher than that for PfHRP2. This underpins the need to further improve the sensitivity of pLDH-based diagnostics.

The bead suspension array described here can successfully be used as for detection and quantification of PfHRP2 and pLDH in whole blood, eluted DBS and plasma or serum samples. The concentration of eluted PfHRP2 from DBS to be equivalent to approximately a 1:20 dilution from whole blood, similarly to previously reported data (40). Differently, for pLDH we found that antigen concentration in eluted DBS corresponds to a 1:60 whole blood dilution, which differs from previously published data showing no differences in antigen recovery between PfHRP2 and pLDH (20). However, such differences could be explained by the different extraction methodologies and storage conditions used.

The quantification of PfHRP2 and pDLH is performed by interpolating MFI values to a regression curve fitted from a calibration curve consisting of recombinant proteins PfHRP2 type A and *P. falciparum* pLDH. Particularly for PfHRP2, the use of a single recombinant protein as a reference material to quantify antigen levels in field samples may provide an approximate estimate of the true concentration. PfHRP2 contains sequences rich in histidine that form the epitopes targeted by the mAbs in RDTs (41), which have been shown to be highly polymorphic in sequence composition of the repeated motifs, as well as in overall length and number of repeated motifs between different parasite strains (41). Baker *et al.* classified PfHRP2 as types A, B, or C depending on the frequency of two epitope repeats (named type 2 and type 7) which confer increased reactivity to mAbs in RDTs (41,42). According to this classification, PfHRP2 Type A comprises the higher number of repeat types 2 and 7, followed by PfHRP2 Type B, and finally PfHRP2 Type C. Our results on the detection of different PfHRP2 types (See Additional file 2: Figure S1A) align with this data and resemble recently published results (24,25).

We observed an overall positive significant correlation between antigen levels and parasite densities similar to what previous studies have found (24), although the correlation was different among the groups of samples analysed (See Additional file 3: Table S1), probably because of the type of sample used for antigen quantification, operational variations and sample storage. Of note, pLDH better correlated with parasite densities compared to PfHRP2. This finding can be explained by the fact that PfHRP2, differently from pLDH, is secreted to the blood stream and persists in circulation for several days. In addition, we observed that the correlation between PfHRP2 and parasite densities was lower in samples from Peru compared to samples from Nigeria, whereas pLDH levels correlated very similarly to parasite densities in both groups of samples. The high number of suspected *P. falciparum* positive samples with *pfhrp2* gene deletions within the group of samples from Peru most probably explains this finding.

## CONCLUSIONS

The quantitative suspension array technology presented here allows for a simultaneous highly sensitive detection of the most commonly used target antigens in malaria RDTs. The assay could be used as a tool to validate next generation RDTs, as well as to estimate malaria burden in endemic areas and to evaluate the impact of malaria control activities. Finally, this assay has the potential to be further upgraded by multiplexing the detection and quantification of antibodies against parasite antigens that could serve as a supplementary tool to study malaria transmission intensity, as well as the detection of other infectious diseases antigens.

## Abbreviations

α: anti
Ag: antigen;
CI: confidence intervals;
DBS: dried blood spot;
EDC: 1-Ethyl-3-[3-dimethylaminopropyl] carbodimide hydrochloride;
ELISA: enzyme-linked immunoabsorbent assay;
GM: Geometric mean;
IgG: immunoglobulin G;
LLOD: lower limit of detection;
LLOQ: lower limit of quantification;
LM: light microscopy;
mAbs: monoclonal antibodies;
MFI: median fluorescence intensity;
p/μl: parasites per microliter;
PCR: polymerase chain reaction;
*Pf*: Plasmodium falciparum;
pg/ml: pictograms per millilitre;
PfHRP2: Plasmodium falciparum histidine-rich protein 2;
PfHRP3: Plasmodium falciparum histidine-rich protein 3;
pLDH: parasite lactate dehydrogenase;
*Pv*: *Plasmodium vivax*;
qPCR: quantitative polymerase chain reaction;
qSAT: quantitative suspension array technology;
RDT: rapid diagnostic test;
RT: room temperature;
SD: standard deviation;
Sulfo-NHS: (N-hydroxysulfosuccinimide);
ULOQ: upper limit of quantification;
µl: microliter;
5PL: five parameters logistic;
°C: degrees Celsius;
%CV: percent coefficient of variation;
%DEV: percent deviation.

## Declarations

### Authors’ contributions

AM and AC conceived and designed the study. CM, EM, ES and RG obtained the plasma samples from Mozambican pregnant women. AV obtained the serum samples from Colombian pregnant women. XD and SD were in charge of studies that allowed collecting the whole blood samples from Senegalese and Peruvian asymptomatic patients, and from Nigerian and Peruvian febrile patients, respectively. AJ and XMV performed all laboratory experiments. AJ, AM and XMV performed the statistical analyses and manuscript preparation. AC, AM, IG and XD provided overall study supervision. All authors read and approved the final manuscript.

## Acknowledgements

We thank the study participants; the staff of the Hospitals, clinical officers, field supervisors and data managers. We acknowledge the teams at CISM in Mozambique, at University of Antioquia in Colombia, at University Cheikh Anta Diop in Senegal, at Universidad Peruana Cayetano Heredia in Peru, and at University of Lagos in Nigeria, who conducted the recruitment of participants and sample processing. We would also like to thank Iveth González, Aida Valmaseda, Marta Vidal and Himanshu Gupta for providing important inputs for optimization of the quantitative bead suspension array; and Laura Puyol, Diana Barrios and Pau Cisteró for providing logistic support. The CISM is supported by the Government of Mozambique and the Spanish Agency for International Development (AECID). ISGlobal is a member of the CERCA Programme, Generalitat de Catalunya.

## Availability of data and material

The datasets used and/or analysed during the current study are available from the corresponding author on reasonable request.

## Consent for publication

Not applicable.

## Ethics approval and consent to participate

The Mozambican National Health and Bioethics Committee, the Medical Research Center Ethics Committee at the Medicine Faculty of Universidad de Antioquia and the Hospital Clinic of Barcelona Ethics Committee approved the use of non-identifiable plasma and serum samples in the current study. Written informed consent was obtained from all participants.

The Senegal National Ethics Committee (Comité National d’Ethique pour la Recherche en Santé) reviewed and approved on 15 January 2015 the study protocol associated with the collection of whole blood specimens from consenting asymptomatic adults in Senegal to support the development and evaluation of new assays for the detection of malaria infections (Protocol SEN14/74). Written informed consent was obtained from all participants.

The Unversidad Peruana Cayetano Heredia Institutioanl Review Board (Comité Institucional de Ética) reviewed and approved on 10 March 2015 the study protocol associated with the collection of whole blood specimens from consenting asymptomatic adults in Peru to support the development and evaluation of new assays for the detection of malaria infections (Protocol 100-02-15). Written informed consent was obtained from all participants.

The study protocol for the evaluation of a multiplex fever diagnostic test was submitted for ethics approval in October 2016 and December 2016 in Peru and Nigeria, respectively, and approvals were obtained in November 2016 in Peru, and January 2017 in Nigeria. Informed consent was obtained from all participants or by legal guardians in cases of underage participants.

The institutional review board at the Universidad Peruana Cayetano Heredia (Lima, Peru) approved the study protocol the specimen collection campaign organized in April 2016 in the area of Iquitos (Peru).

## ADDITIONAL FILES

### Additional file 1

#### Format

docx format.

#### Tittle of data

<u>Supplementary Materials and Methods</u>

#### Data

Includes “Text S1. Biotinylation of detection mAbs”, “Text S2. Bead suspension array procedure”, “Text S4. Singleplex versus Multiplex testing”, and “Text S4. Preparation and extraction of proteins from dried blood spots”

### Additional file 2

#### Format

pdf format

#### Tittle of data

Supplementary figure 1.

#### Description of data

**Figure S1. Assay optimization.** (**A**) Serial dilutions of recombinant PfHRP2 types A, B and C were assayed to determine the lowest concentration at which each antigen is detected; (**B**) *P. falciparum* Benin I and Borneo and *P. vivax* field isolates were assayed in a serial dilution fashion to assess differences between the analytical sensitivity for *P. falciparum* and *P. vivax* pLDH; (**C**) PfHRP2 and pLDH positive samples (plasma, cultured field isolates and recombinant proteins) were assayed in singleplex (X axes) and multiplex (Y axes).

### Additional file 3

#### Format

docx format

#### Tittle of data

Supplementary Tables 1 and 2.

#### Description of data

Table S1. Correlation between antigen levels and parasite densities for each group of clinical samples analysed.

