## Additional file 1_Texts S1, S2, S3 and S4 for "Quantification of malaria antigens PfHRP2 and pLDH by quantitative suspension array technology in whole blood, dried blood spot and plasma"

**Supplementary Material and methods**

**Text S1. Biotinylation of detection mAbs.** Biotinylation of α-PfHRP2 (MBS834434, MyBioSource, San Diego, CL) and α-PAN-pLDH (PA-2, AccessBio, Somerset, NJ) antibodies was performed following manufacturer instructions. However, upon biotinylation, mAbs did not provide the expected the fluorescence range offered by the Luminex technology. To achieve higher MFI values, biotinylated mAbs underwent a second round of biotinylation, following again the manufacturer instructions. After each round of biotinylatioon, the antibody concentration in the collected flow-through was measured by spectrophotometry (Epoch Microplate Spectrophotometer, BioTek). The antibody solution was adjusted to the desired concentration, aliquoted and stored at 4 °C.

**Text S2. Bead suspension array.** Biological samples were incubated overnight at 4 °C in agitation (500 rpm) with 2000 magnetic beads per analyte in 96-well flat bottom plate. The plate was washed by pelleting microspheres using a magnetic separator (40-285, EMDMillipore, Burlington, MA) and re-suspended with wash buffer (0.05% Tween 20/PBS). 100 µL of biotinylated antibodies α-HRP2 and α-PAN-pLDH at 1 µg/ml was applied to all wells and incubated at RT for 2 hours with agitation and protection from light. The beads were then washed again and incubated with streptavidin-PE (42250-1ML, Sigma Aldrich, St. Louis, MO) diluted 1:1000 in assay buffer for 30 minutes in gentle agitation in the dark. Finally, the beads were washed and resuspended in assay buffer, and the plate was read using the Luminex xMAP® 100/200 analyser (Luminex Corp., Austin, TX). A minimum of 50 microspheres per analyte were acquired per spectral signature and results were exported as crude median fluorescent intensity (MFI). Background (blank) MFIs were subtracted and normalized (nMFI) to account for plate to plate variation. Normalization was done by dividing the blank subtracted MFI of each sample by the MFI value corresponding to an interpolated arbitrary concentration to the standard curve of the same plate, and then multiplied by the mean arbitrary concentration MFI value. Quantification was performed against a 5-parameter logistic (5-PL) regression curve with logarithmic variance weighting fitted from a calibration curve consisting of recombinant proteins PfHRP2 type A and Pf pLDH.

**Text S3. Singleplex *versus* Multiplex testing.** Multiplexed immunoassays can present cross-reactions between protein antigens and antibodies, leading to inaccurate results and erroneous conclusions. Serial dilutions of recombinant PfHRP2 type A and *Pf* pLDH proteins were tested in singleplex and multiplex formats, and resulting fluorescence signals were compared to assure no cross-reactivity was occurring. In addition, a selection of *P. falciparum* cultured field strains (CDC Benin, CDC Borneo, CDC Santa Lucia) and a pool of *P. falciparum* positive plasma samples at different dilutions were also tested in both formats.

**Text S4. Extraction of proteins from dried blood spots.** DBS were prepared by spotting 25 μl onto Whatman 3 filter papers (W3). Papers were dried overnight at RT, wrapped in aluminium foil and stored at -20 °C in sealing bags with silica gel. Extraction of blood proteins was performed on three 3 mm diameter punches. DBS were cut using a manual punch (WHATWB100078, Uni-CoreTM 3MM, GE Healthcare, Chicago, IL) and placed in 1.5 ml microcentrifuge tubes with 150 μl assay buffer. The tubes were agitated at 1000 rpm overnight at room temperature. The day after, tubes were centrifuged to spin down the filter paper, supernatants were transferred to 0.5 ml cryopreservation tubes and stored at -80 °C. The dilution applied to PfHRP2 and pLDH at the time of eluting blood from DBS was calculated by dividing the antigen concentration in the original whole blood sample and the concentration in the corresponding eluted DBS solution, and then multiplied by 100:

*Dilution = (antigen concentration in WB/antigen concentration in eluted DBS solution)*100*
