## Supplementary figures and images for "Quantification of malaria antigens PfHRP2 and pLDH by quantitative suspension array technology in whole blood, dried blood spot and plasma"

### Additional file 2_Figure S1

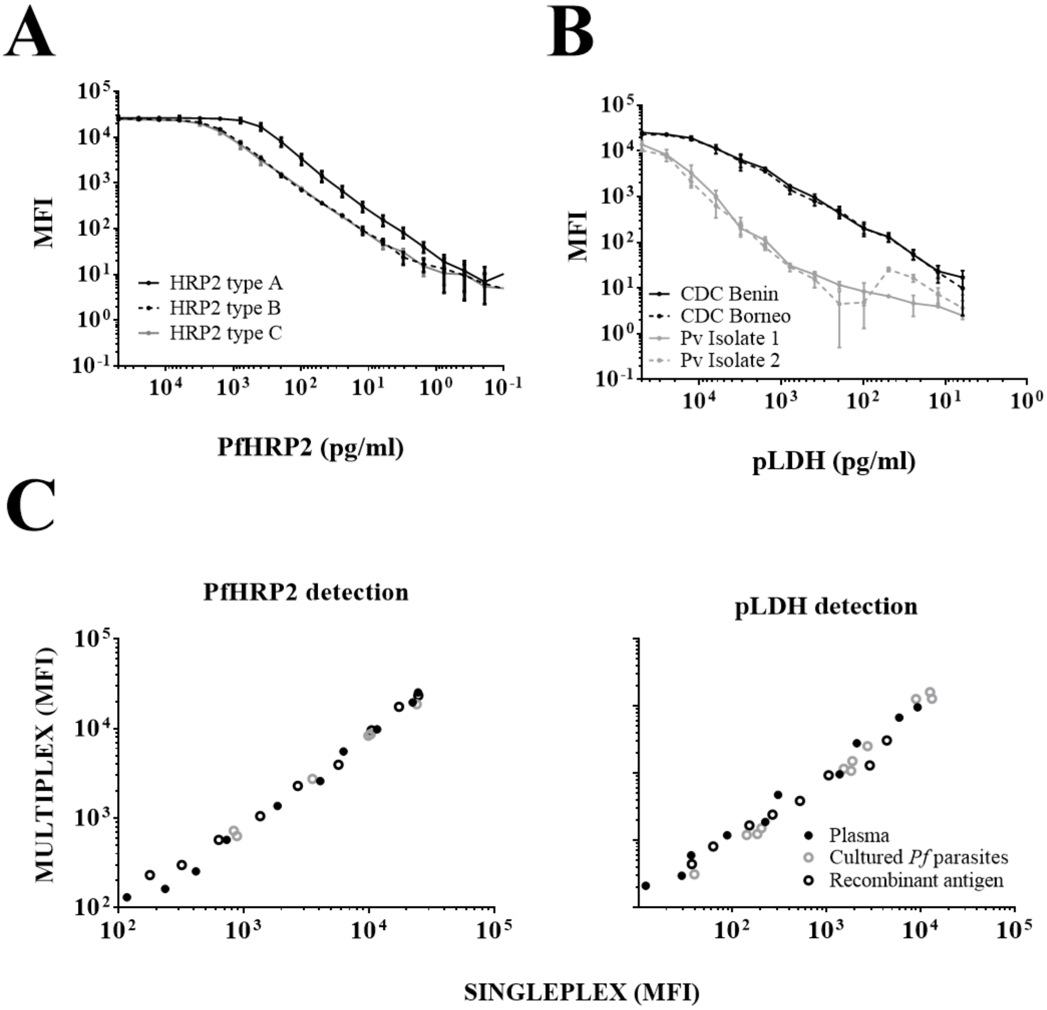
