## Additional file 3_Table S1 for "Quantification of malaria antigens PfHRP2 and pLDH by quantitative suspension array technology in whole blood, dried blood spot and plasma"

**Table S1. Correlation between antigen levels and parasite densities for each group of clinical samples analysed.**

|  |  |  | **PfHRP2** | | | **pLDH** | | |
| --- | --- | --- | --- | --- | --- | --- | --- | --- |
| **Sample origin** | **Sample type** | **Method** | **Number of pairs** | **Spearman r**  **(95% CI)** | **P value** | **Number of pairs** | **Spearman r**  **(95% CI)** | **P value** |
| Senegal | Whole blood | PCR | 39 | 0.55 (0.27-0.74) | 0.0003 | 26 | 0.84 (0.66-0.93) | < 0.0001 |
| Peru | Whole blood | PCR | NA | NA | NA | 5 | 0.30 (-) | 0.683 |
| Mozambique | Plasma | PCR | 30 | 0.80 (0.61-0.90) | < 0.0001 | 16 | 0.91 (0.74-0.97) | < 0.0001 |
| Colombia | Serum | Microscopy | 23 | 0.18 (-0.26-0.56) | 0.414 | 19 | 0.10 (-0.38-0.54) | 0.679 |
| Nigeria | Whole blood | Microscopy | 243 | 0.47 (0.36-0.56) | < 0.0001 | 241 | 0.78 (0.72-0.82) | < 0.0001 |
| Peru | Whole blood | Microscopy | 120 | 0.20 (0.01-0.37) | 0.0308 | 187 | 0.76 (0.70-0.82) | < 0.0001 |
| ALL |  |  | 457 | 0.45 | < 0.0001 | 494 | 0.76 | < 0.0001 |
| NA: Not applicable | | | | | | | | |
